# SMAD4: A Critical Regulator of Cardiac Neural Crest Cell Fate and Vascular Smooth Muscle Differentiation

**DOI:** 10.1101/2023.03.14.532676

**Authors:** Brianna E. Alexander, Huaning Zhao, Sophie Astrof

## Abstract

**Background:** The pharyngeal arch arteries (PAAs) are precursor vessels which remodel into the aortic arch arteries (AAAs) during embryonic cardiovascular development. Cardiac neural crest cells (NCs) populate the PAAs and differentiate into vascular smooth muscle cells (vSMCs), which is critical for successful PAA-to-AAA remodeling. SMAD4, the central mediator of canonical TGFβ signaling, has been implicated in NC-to-vSMC differentiation; however, its distinct roles in vSMC differentiation and NC survival are unclear.

**Results:** Here, we investigated the role of SMAD4 in cardiac NC differentiation to vSMCs using lineage-specific inducible mouse strains in an attempt to avoid early embryonic lethality and NC cell death. We found that with global SMAD4 loss, its role in smooth muscle differentiation could be uncoupled from its role in the survival of the cardiac NC *in vivo*. Moreover, we found that SMAD4 may regulate the induction of fibronectin, a known mediator of NC-to-vSMC differentiation. Finally, we found that SMAD4 is required in NCs cell-autonomously for NC-to-vSMC differentiation and for NC contribution to and persistence in the pharyngeal arch mesenchyme.

**Conclusions:** Overall, this study demonstrates the critical role of SMAD4 in the survival of cardiac NCs, their differentiation to vSMCs, and their contribution to the developing pharyngeal arches.

## 1. Introduction

The cardiovascular system is the first organ system formed during embryonic development ^1,2^. Defects affecting the heart and/or its associated vasculature are known as Congenital Heart Disease (CHD).

CHD is one of the most common human birth defects, affecting ∼1% of all live births annually ^3^. It is also the number one cause of infant mortality, with an estimated 25% of all CHD cases so severe, termed “critical CHDs,” that they necessitate immediate surgical intervention ^4–6^. Longitudinal studies revealed that while 82.5% of all critical CHD patients survive to 1 year, only 68.8% survive to adulthood (18 years of age) ^4^. Despite these staggering statistics, only a reported 15% of all CHD cases are linked to a known cause, underscoring the need to elucidate the etiology of CHD so that new preventative and treatment options can be developed ^7^.

Critical CHDs encompass phenotypes affecting the aortic arch arteries (AAAs), the vessels which connect to the aortic arch and route oxygenated blood throughout systemic circulation. Any occlusion or malformation of these vessels disrupts blood flow throughout the body and often requires immediate surgical intervention for survival ^8,9^. During cardiovascular development, the AAAs originate from three pairs of symmetrical vessels, the 3^rd^, 4^th,^ and 6^th^ pharyngeal arch arteries (PAAs), which undergo remodeling between E10.5-E13.5 in murine embryos to form the AAA tree ^10,11^. PAA-to-AAA remodeling is predominantly facilitated by the ectodermal-derived cardiac neural crest cell (NC) population. Cardiac neural crest cells (NCs) belong to a subset of cranial NCs. They migrate towards the embryonic outflow tract and contribute to forming the aorticopulmonary septum. This structure helps to separate the pulmonary trunk and aorta. Additionally, cardiac NCs migrate to the posterior pharyngeal arches (3-6) to aid in the development of the aortic arch arteries ^12,13^.

When NCs migrate into the 3^rd^-6^th^ pharyngeal arches, they form layers around the endothelium of the PAAs. NCs closest to the endothelium differentiate into vascular smooth muscle cells (vSMCs), thereby contributing to the tunica media ^14,15^. This NC-to-vSMC differentiation is critical to proper AAA morphogenesis as it enables the developing vessel to maintain physical integrity under the forces of blood flow and vascular remodeling ^13,16,17^. Processes that interfere with normal NC activity in the pharyngeal region, including defective NC migration, vSMC differentiation, or NC survival, can result in arch artery regression and severe AAA defects. An example of this is interrupted aortic arch (IAA), a lethal phenotype caused by either defective formation or premature regression of the 4^th^ left PAA, which is seen in 50-89% of 22q11.2 patients ^8,18–20^. Given the severity of 4^th^ PAA-related phenotypes, the 4^th^ PAAs in particular, were analyzed in this study.

Several genetic mutations have been implicated in CHD cardiovascular phenotypes, including *JAG1*, linked to Alagille syndrome, *CHD7*, linked to CHARGE syndrome, *TBX1*, linked to 22q11.2 syndrome, ^21^ and recently SMAD4, linked to Myhre syndrome ^22,23^. SMAD4 is the common signaling component in the canonical TGFβ and BMP signaling pathways. In addition to its role as a tumor suppressor ^24,25^, SMAD4 is known to be critical for proper gastrulation and embryonic development, with global mutants resulting in mouse embryonic lethality by E8.5 due to improper development of the mesoderm and visceral endoderm (Sirard et al., 1998; Yang et al., 1998). In recent years, SMAD4 mutations have been linked to various cardiovascular phenotypes, including coarctation of the aorta, ventricular septal defects, atrial septal defects, and persistent truncus arteriosus ^22,23^. Recently, the SMAD4 Tyr95 mutation has been identified as a happloinsufficient mutation which increases patient susceptibility to CHD ^26^. Patients presented with defects including ventricular septal defects, bicuspid aortic valve, thoracic aortic aneurism, and patent ductus arteriosus ^26^. Although SMAD4 has emerged as a factor in CHD etiology, its role in NC contribution to aortic arch artery morphogenesis is not well understood.

Many groups have investigated the role of TGFβ and SMAD4 in the neural crest during cardiovascular morphogenesis using the Wnt1-Cre1 transgenic mouse strain ^27^. Through Wnt1-Cre1-mediated deletion of SMAD4, TGFβ receptor 1 (Alk5), TGFβ receptor 2, BMP receptor 1 (Alk2) ^28–31^, and over-expression of the TGFβ inhibitor, Smad7 ^32^, it has been reported that TGFβ/SMAD4 is important for the survival of cardiac NCs that contribute to the cardiac outflow tract and pharyngeal arches. However, it remains unclear whether the role of SMAD4 in NC differentiation to vSMCs *in vivo* is independent of its role in NC survival. In addition, the use of the Wnt1-Cre1 transgenic mouse model may complicate the interpretation of results. In this strain, Wnt1 and Wnt1 signaling are ectopically activated due to the features of the transgene ^33^. Wnt1, along with BMPs and FGFs, regulates NC induction and thus, if over-expressed, may potentially rescue phenotypes in NC genetic mutants ^34–36^. Although it is unknown whether NCs in the Wnt1-Cre1 strain overexpresses Wnt1, phenotypic disparities between knockout phenotypes using Wnt1-Cre1 and other NC Cre drivers have been reported ^35–38^. Given the critical roles of SMAD4 in various tissues, determining if vSMC defects are a consequence of cell death or if vSMC differentiation can be regulated independently of cell death would help to clarify the role of SMAD4 in CHD etiologies.

To investigate the specific role of SMAD4 in the cardiac NC and to distinguish its role in NC differentiation from its role in NC survival, we generated conditional knockout mouse models to ablate SMAD4 in a tissue-specific and temporal manner. We examined the expression of αSMA (as a marker of vSMCs), and the extracellular matrix protein, Fibronectin (Fn1), a known regulator of NC-to-vSMC differentiation ^37^, as a potential mechanism for SMAD4 regulation of vSMC differentiation. Our findings suggest that in global SMAD4 mutants, SMAD4 regulation of αSMA can be separated from its role in NC survival *in vivo*, and that SMAD4 may be a potential regulator of Fn1 induction *in vivo*. Additionally, we demonstrate that while SMAD4 expression in the endothelium is not required for Fn1 or αSMA expression, that SMAD4 is required, in a cell-autonomous manner, for cardiac NCs to contribute to and persist in the developing pharyngeal arches. Without SMAD4, cardiac NCs do not persist in the pharyngeal arches. This study highlights the requisite role of SMAD4 in the development of the cardiac NC and its contribution to the pharyngeal arch mesenchyme.

## 2. Results

### 2.1 SMAD4 regulates smooth muscle differentiation around the 4^th^ pharyngeal arch arteries

To examine the role of SMAD4 in αSMA expression and Fn1 induction in the PAAs, we assayed the expression of each in the pharyngeal arches of E12.5 embryos. In E12.5 embryos, the process of PAA-to-AAA remodeling has begun, as demonstrated from the regressing right 6^th^ PAA **(Fig. 1A-B)** and *SMAD4 mRNA* is ubiquitously expressed in the pharyngeal mesenchyme **(Fig. 1C-C’)**. *Fn1 mRNA* and protein are enriched around the PAA endothelium **(Fig. 1D-E’)**, and cardiac NCs have differentiated to vSMCs **(Fig. 1F-F’)**. Fn1 is a known mediator of NC-to-vSMC differentiation, evidenced by its robust expression and co-localization with αSMA around the PAAs **(Fig. 1E-F’)** ^37^.

**Figure 1.**
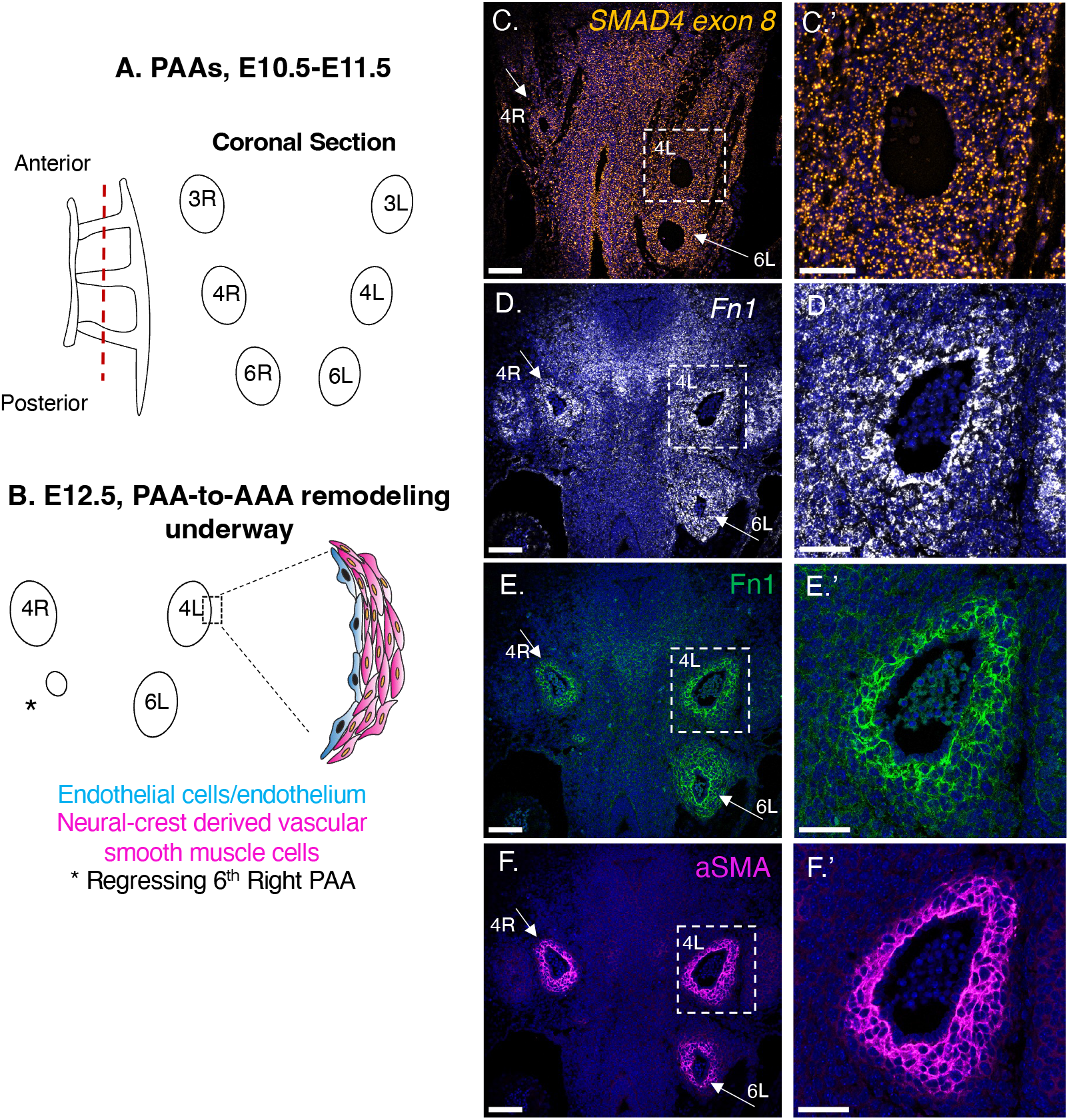
SMAD4 and Fn1 are highly expressed in the pharyngeal arches at E12.5. **A-B.** Schematic depiction of PAAs at E10.5 – E12.5. The paired PAAs, 3rd,4th and 6th (**A**) are remodeled between E11.5-E13.5 (**B**). Dashed red line in **A** indicates the plane of coronal section. Coronal sections through E12.5 PAs in a control embryo show expression of *SMAD4* mRNA (**C-C’**), *Fn1* mRNA (**D-D’**), Fn1 protein (**E-E’**) and αSMA (**F-F’**). White arrows point to the 4th right and 6th PAA. The 4th left pharyngeal arch (boxed) is magnified in **C’-F’**. Scale bars in **C-F** are each 100 µm. Scale bars in **C’-F’** are 50 µm.

The ablation of SMAD4 during early development results in embryonic lethality ^39,40^ and causes NC cell death ^41,42^. To avoid early embryonic lethality and NC death and to test the later roles of SMAD4 in the differentiation of NC into vSMCs, we sought to induce the deletion of *SMAD4* in a narrow window of time by using the R26R^CreERT2/CreERT2^ mouse strain ^43^, in which Cre expression can be induced ubiquitously. The 4^th^ pharyngeal arches form during the embryonic day E10.5 of development ^44^. In the morning of E10.5, there is a vascular plexus, and by the evening, the patent arch artery is formed, with NC-vSMC differentiation to follow at E11.5. Thus, to globally ablate SMAD4 before NCs differentiate into vSMCs around the 4^th^ PAA, and to avoid lethality due to SMAD4 deletion, we injected tamoxifen at 8 am on day E10.5 of development **(Fig. 2A)**. Since Fn1 is highly induced in NCs around the PAAs at E11.5, concomitant with the onset of NC-to-vSMC differentiation ^37^, embryos were analyzed at E12.5 to examine the effect of SMAD4 loss on the expression of αSMA and Fn1, and to allow adequate time for Cre-mediated loss of SMAD4.

**Figure 2.**
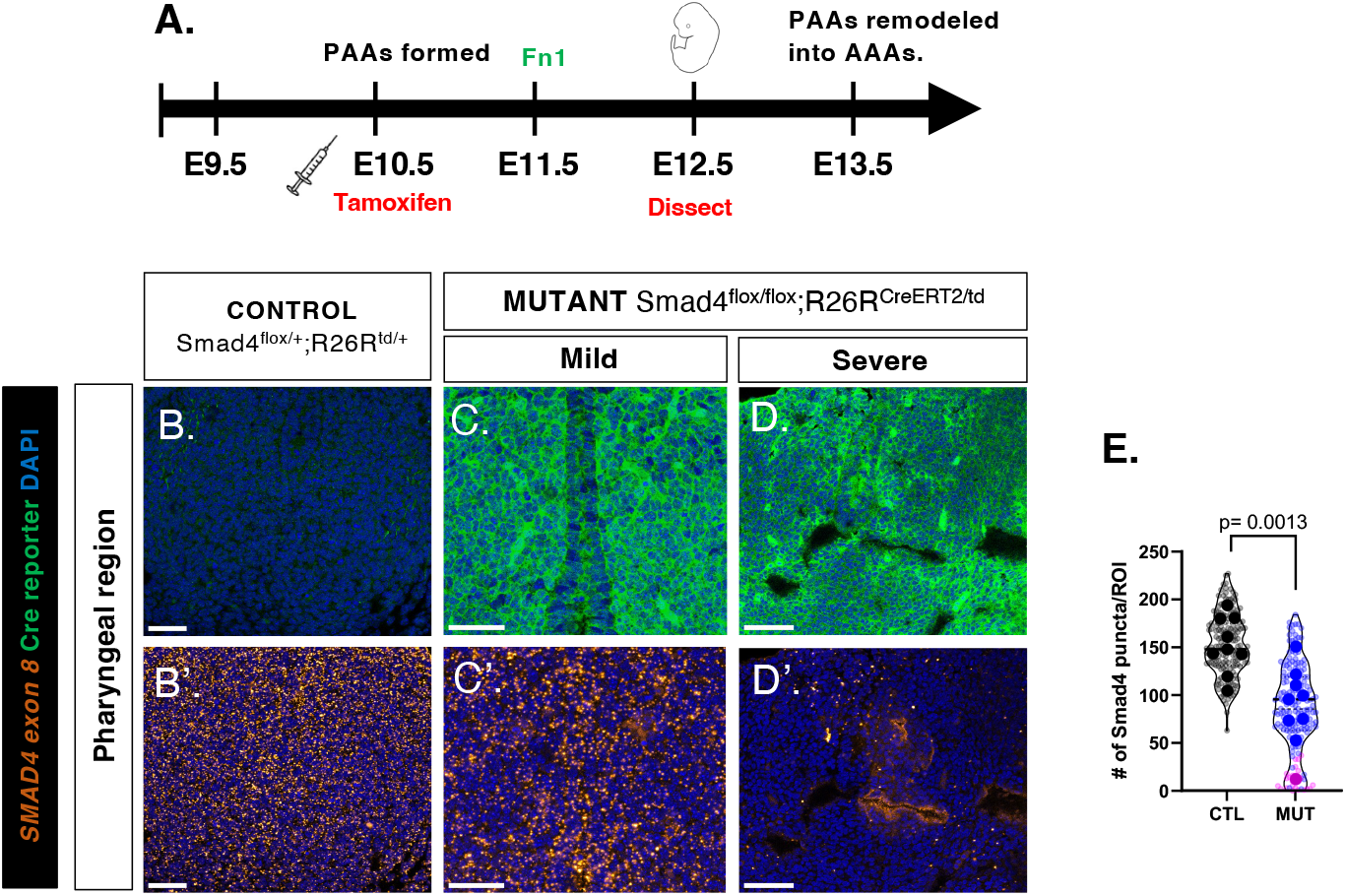
R26R^CreERT2^-mediated recombination results in graded *SMAD4* knockdown. Tamoxifen injection conditions (**A**). Images of *SMAD4* exon 8 fluorescent *in-situ hybridization* and immunofluorescence for the Cre reporter in the pharyngeal region of a control (**B-B’**), mild mutant (**C-C’**) and the severe mutant (**D-D’**) at E12.5. *SMAD4* exon 8 mRNA quantified in (**E**), with the severe mutant (N=1, pink) having fewer than 50 puncta per area and mild mutants (N=8, blue) having more than 50 puncta per area. Large dots are embryo average and small dots are individual sections analyzed. 2-tailed, unpaired, Student’s t-test was used. N= 9 controls, 9 mutants; p <0.05 considered statistically significant. Scale bars are 50 µm.

Cre-mediated deletion of *SMAD4* exon 8 in the SMAD4^flox/flox^ strain results in a SMAD4-null allele (Yang et al., 2002). To select embryos for further molecular and phenotypic analyses, we initially analyzed SMAD4 ablation by performing Western blot to assay the expression of SMAD4 protein in the posterior region of each embryo **(Supp. Fig. 1A-C)**. 47 Smad4^flox/flox^;R26R^CreERT2^ mutant embryos were analyzed, and 9 embryos exhibiting, on average, a ∼5-fold decrease in SMAD4 protein expression compared to Cre-negative controls were selected for further analyses. Mendelian ratios for each genotype were observed **(Supp. Table 1)**. To evaluate the efficiency of Cre-mediated recombination in the pharyngeal region, we first assayed the expression of the tdTomato Cre reporter, which revealed efficient recombination in the vast majority of cells after tamoxifen injection **(Fig. 2B-D)**. We then assayed *SMAD4 exon 8 mRNA* in the pharyngeal region by *in situ* hybridization. Sections were analyzed by counting the number of *SMAD4 exon 8-*containing mRNA puncta within a defined area. Compared with controls, which contained ∼100-200 SMAD4+ puncta per ROI, these analyses revealed two classes of mutant embryos: mild, with more than 50 puncta per area, and severe, with less than 50 puncta **(Fig. 2B’-D’)**. Of 9 mutant embryos, 8 were in the mild category based on *SMAD4* mRNA expression, and 1 mutant embryo fell into the severe class **(Fig. 2E,** pink). Together, these analyses demonstrate that although Cre-mediated recombination was efficient at the reporter locus, the loss of SMAD4 expression was varied.

To assay the consequences of SMAD4 deletion on vSMC differentiation and Fn1 expression, we further analyzed the 9 mutant embryos previously selected at E12.5. At E12.5, αSMA was expressed in multiple cell layers around the pharyngeal arch of the control embryo **(Fig. 3A-A’)**. Out of 9 mutants, 6 had attenuated αSMA expression levels, including the severe mutant **(Fig. 3B-B’, C-C’)**, while three exhibited αSMA levels similar to controls **(Fig. 3D-D’)**, quantified in **(Fig. 3E-F)**. To determine if the reduction in αSMA was a result of changes in NC proliferation or survival, TUNEL and pHH3 staining were performed on consecutive sections through the pharyngeal arches **(Fig. 3G-I’)**. The total cell number and cell proliferation were not affected in the mutants (**Fig. 3J, K**). However, although rare, we found a statistically significant increase (from 1.2% to 2.5%) in the fraction of apoptotic cells in mutant embryos **(Fig. 3L)**. Since the majority (>97%) of cells surrounding the 4^th^ PAA were TUNEL-negative, the decreased NC-to-vSMC differentiation seen in 60% of the mutants cannot be attributed to cell death. To determine if SMAD4 regulated the expression of Fn1, we then examined *Fn1 mRNA* and protein at E12.5. We did not observe any differences between controls and mild mutants **(Fig. 4A-B’)**. However, the severe mutant, which had the highest degree of SMAD4 ablation, showed a significant attenuation of *Fn1 mRNA* and protein **(Fig. 4C-C’)**, quantified in **(Fig. 4D-E)**. Taken together, these data indicate that SMAD4 regulates αSMA expression independently of survival and suggests that the expression of Fn1 may also depend on SMAD4 levels.

**Figure 3.**
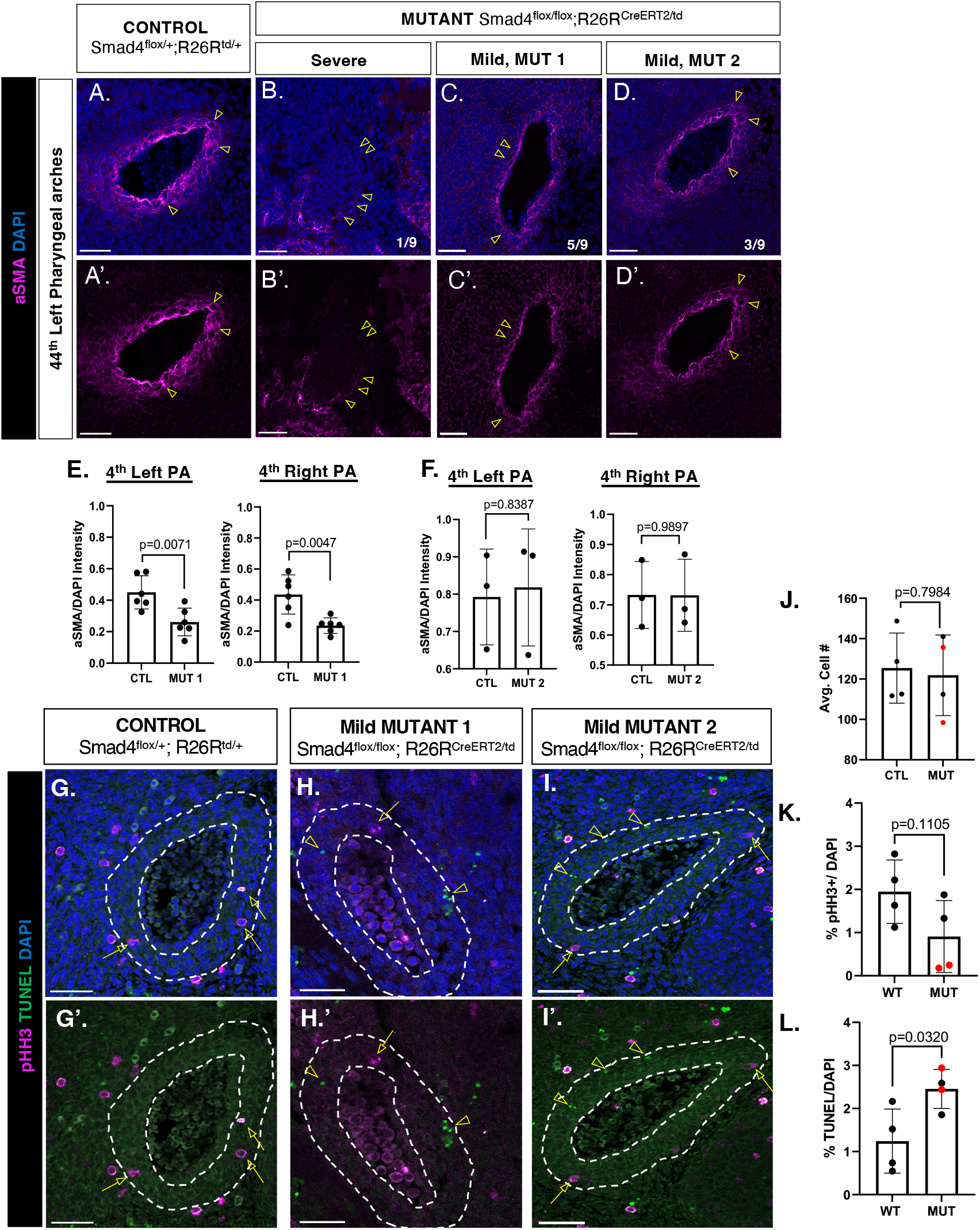
*SMAD4* regulates smooth muscle differentiation. Representative images of αASMA expression around the 4th PA of a control (**A-A’**), the severe mutant (**B-B’**), and two classes of mild mutants (**C-D’**). Yellow arrowheads point to αSMA expression. Quantifications of αSMA expression (**E-F**). N=9 controls and 9 mutants analyzed. TUNEL and pHH3 staining were performed on consecutive control (**G-G’**) and mutant (**H-I’**) sections; arrows point to pHH3+ cells and arrowheads point to TUNEL+ cells. (**J-L**) Quantifications of % pHH3+ and % TUNEL+ cells in 4-cell layers surrounding the PAA endothelium outlined by the white dashed lines in (**G-I’)**. N=4 controls and N=4 mutants, affected (red) and unaffected mutants (black) are marked. Each dot is an average of all sections analyzed per embryo. 2-tailed, unpaired Student’s t-test was used for statistical analyses. Scale bars in **A-C’** are 50 µm, **D-D’** where scale bars are 35 µm. Scale bars in **G-G’** are 42 µm, in **H-H’** are 50 µm and in **I-I’** are 30 µm.

**Figure 4.**
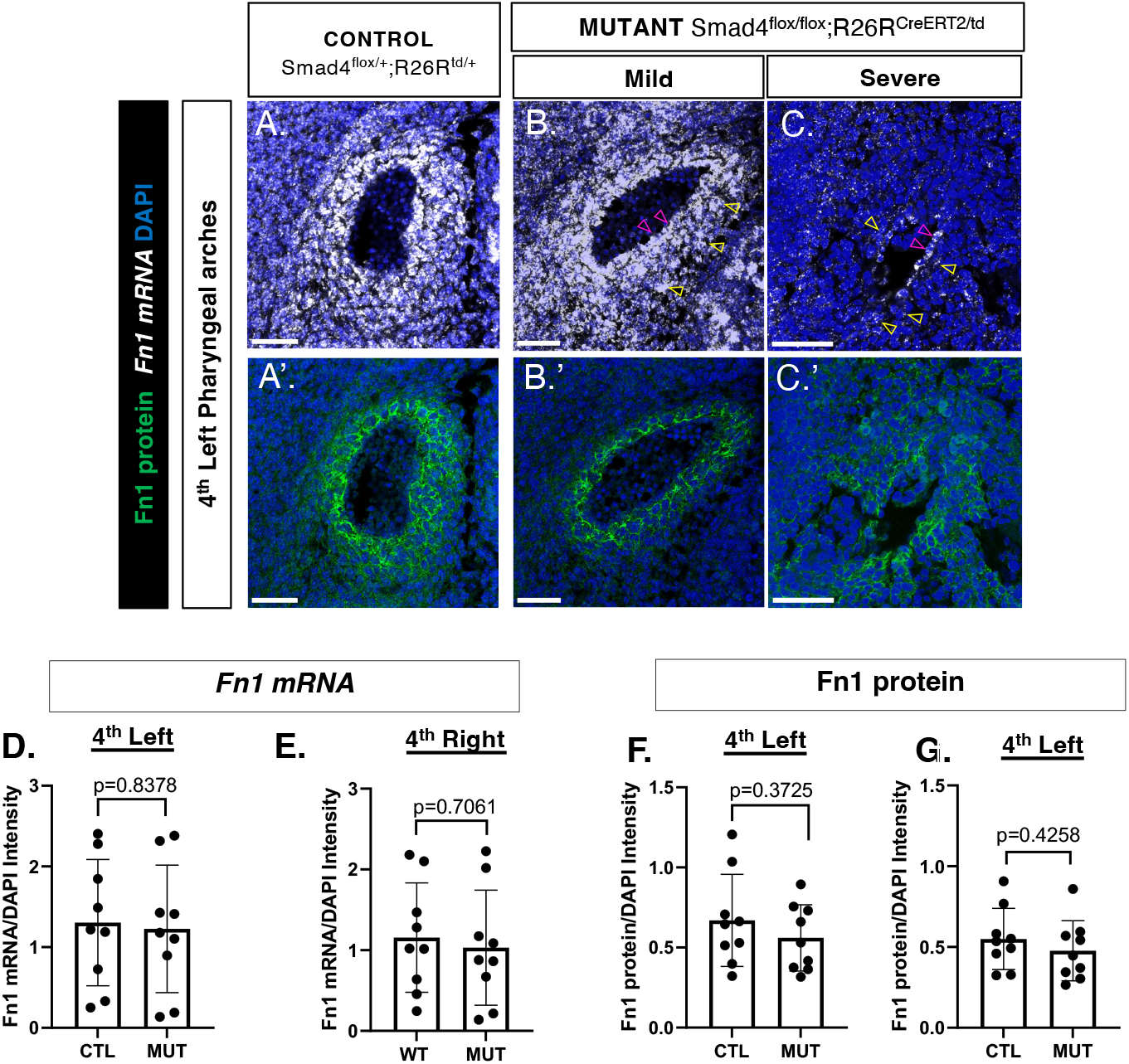
Global loss of SMAD4 results in attenuated Fn1 expression around the 4^th^ PAA in the severe Smad4^flox/flox^; Rosa^CreERT2^ mutant. Images of *Fn1 mRNA* and immunofluorescence for Fn1 protein on coronal sections through the 4^th^ PAs in an E12.5 control (**A-A’**), mild mutant (**B-B’**) and severe mutant (**C-C’**). Yellow arrowheads point to *Fn1 mRNA* in NC cells around the endothelium and pink arrowheads point to *Fn1 mRNA* in endothelial cells. Quantifications of *Fn1 mRNA* are shown in (**D-E**) and Fn1 protein shown in (**F-G**). N= 9 controls and N= 9 mutants. Each dot is an average of all sections analyzed per embryo. For statistics, 2-tailed, unpaired, Student’s t-test was used; p <0.05 is considered statistically significant. Scale bars are 50 µm.

### 2.2 The expression of SMAD4 in the PAA endothelium is not required for Fn1 or αSMA expression

Although only 1 severe SMAD4 mutant was collected, our results suggested that SMAD4 was important for both αSMA expression and, possibly, for Fn1 induction in NCs within the pharyngeal arches. As we could not obtain more viable severe SMAD4 mutants, our focus shifted to identifying the specific cell type that required SMAD4 for inducing Fn1 and αSMA expression. In addition, we hypothesized that targeting SMAD4 in a tissue-specific manner, instead of globally, would result in more efficient deletion of SMAD4 and prevent embryonic lethality.

The PAA endothelium is directly adjacent to NCs **(Fig 1B)**. A previous conditional knockout study using *Tie2-Cre* revealed that SMAD4 loss in endothelial cells resulted in embryonic lethality by E10.5, with mutant embryos exhibiting cardiovascular defects, aberrant vascular patterning, and abnormal deposition of the ECM protein, laminin, around the dorsal aortae ^46^. This study suggested a role for endothelial SMAD4 in cardiovascular morphogenesis; however, the earlier embryonic lethality in this strain prevented the analysis of later defects.

To ablate SMAD4 in the endothelium and to avoid embryonic lethality, we used the Cdh5-CreERT2 inducible mouse strain (Dr. Ralf Adams, Max Planck Institute for Molecular Biomedicine) and injected tamoxifen at E10.5 **(Fig. 5A)**. We then assayed the efficiency of SMAD4 deletion in E12.5 embryos. We found that the ablation of SMAD4 in the endothelium of Smad4^flox/flox^;Cdh5-CreERT2 mutants was highly efficient, with an average of 76% of Cre-reporter+ cells lacking *SMAD4* in five mutants from three independent experiments **(Fig. 5B-E”)**.

**Figure 5.**
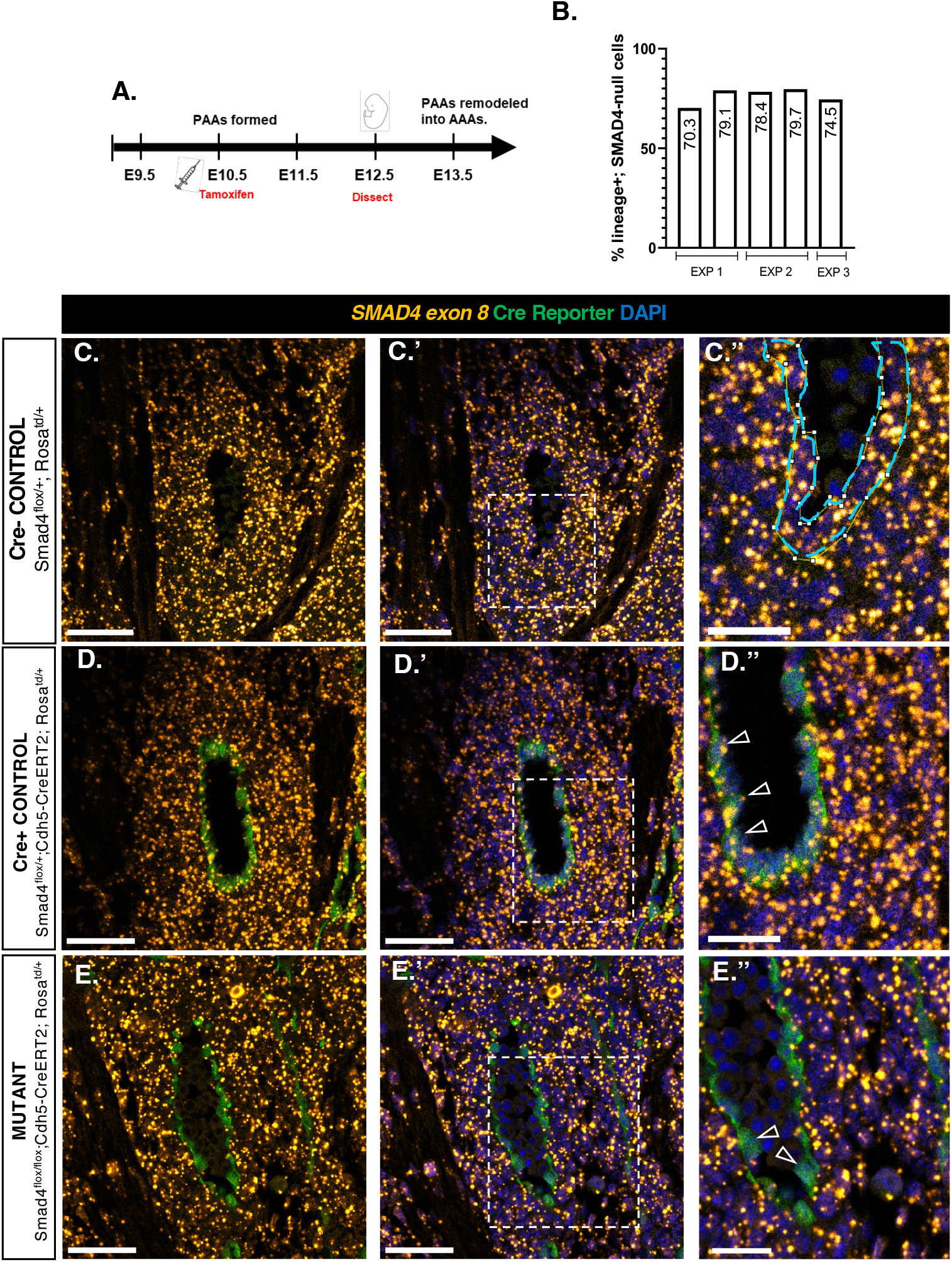
Cdh5-CreERT2-mediated SMAD4 loss is efficient in the PAA endothelium. Tamoxifen injection conditions (**A**). Percentage of Cdh5-lineage+; SMAD4-null cells quantified in the endothelium of five E12.5 mutants from N=3 individual experiments (**B**). Representative images of fluorescent *in situ hybridization* for SMAD4 exon 8 and immunofluorescence for the Cre reporter in a Cre negative control (**C-C’’**), Cre positive control (**D-D’’**) and mutant (**E-E’’**). Arrows point to Cre+; SMAD4+ cells in **D’’** and Cre+; SMAD4-null cells in **E’’**. Scale bars in **C,-C’, D-D’** and **E-E’** are 50 µm and scale bars in **C’’, D’’** and **E’’** are 25 µm.

Quantification of Fn1 mRNA and protein in E12.5 Smad4^flox/flox^;Cdh5-CreERT2 mutants demonstrated that the loss of SMAD4 from the endothelium did not affect *Fn1 mRNA* or protein in NC-derived cells surrounding the 4^th^ PAAs **(Fig. 6A-A’, B-B,’ and C-F)**. The differentiation of NCs to αSMA+ vSMCs was also unaffected **(Fig. 6A’’, B’’ and G-H)**. Additionally, the loss of SMAD4 from the endothelium did not perturb *Fn1 mRNA* expression in the endothelium itself **(Fig. 6I-L)**. These findings indicate that endothelial SMAD4 is not required for Fn1 expression in the NC-derived cells surrounding the 4^th^ PAAs, or their subsequent differentiation into vSMCs. Therefore, these studies suggested that the expression of SMAD4 was required by NCs for NC-to-vSMC differentiation and morphogenesis of the aortic arch arteries.

**Figure 6.**
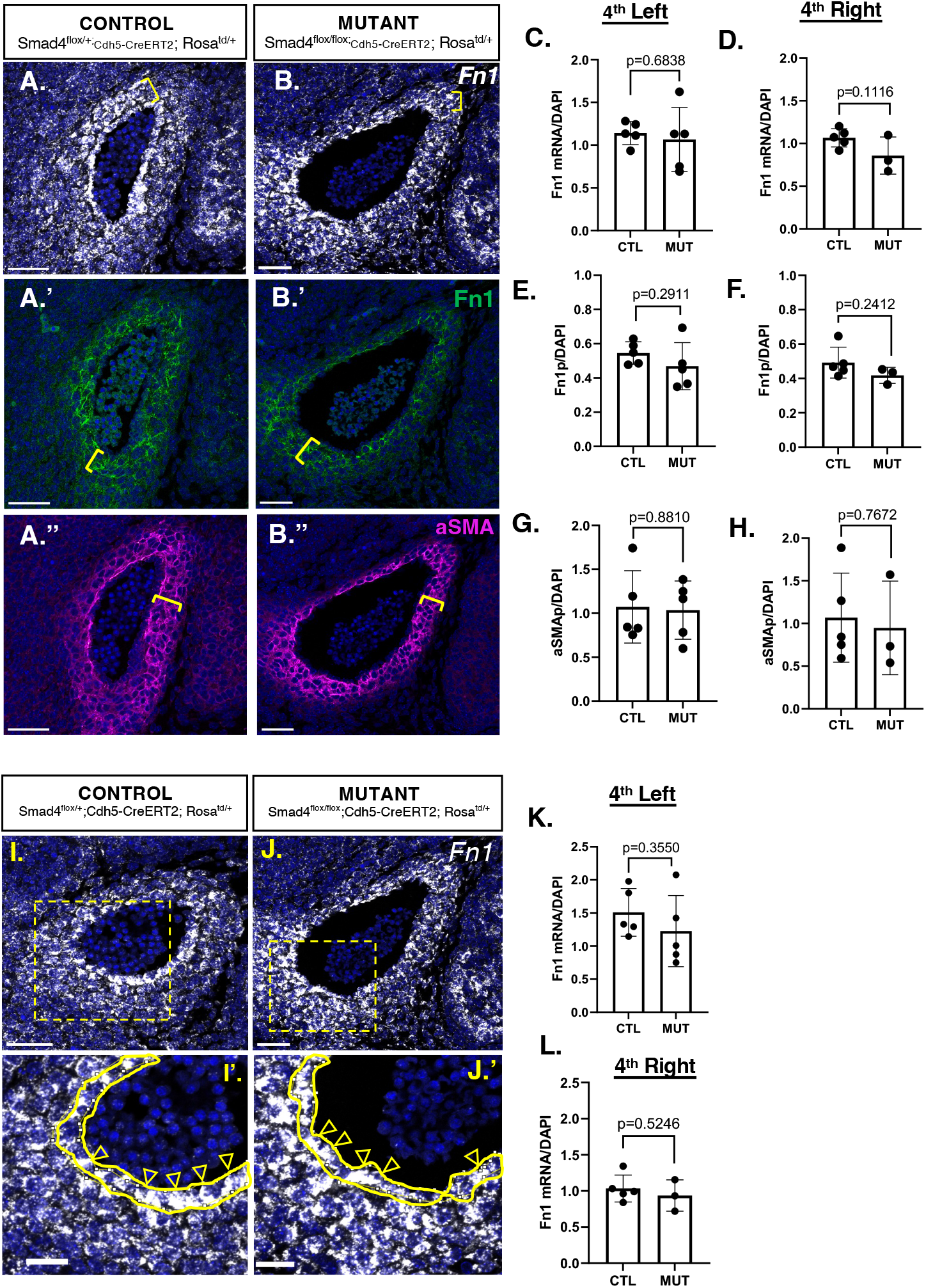
Endothelial loss of SMAD4 does not attenuate Fn1 or αSMA in the neural crest. Images of fluorescent *in situ* hybridization for *Fn1* mRNA, and immunofluorescence for Fn1 protein, and αSMA in the 4th PA of an E12.5 control (**A-A’’**) and mutant (**B-B’’**) embryo. Neural crest cell layers closest to the endothelium are indicated by yellow brackets. Quantifications of *Fn1* mRNA (**C-D**), Fn1 protein (**E-F**) and αSMA (**G-H**). Images of *Fn1* mRNA in control (**I-I’**) and mutant (**J-J’**). Open arrowheads indicate *Fn1* mRNA in the endothelium. Quantifications of *Fn1* mRNA in the endothelium (**K-L**). N= 5 controls and 5 mutants. Each dot is an average of all sections analyzed per embryo. For statistics, 2-tailed, unpaired, Student’s t-test was used; p <0.05 is considered statistically significant. Scale bars in **A-J** are 50 µm and scale bars in **I’** and **J’** are 25 µm.

### 2.3 The expression of SMAD4 in NC-derived cells is required for their contribution to the pharyngeal mesenchyme

Cardiac NC-derived cells comprise multiple cell layers immediately adjacent to the PAA endothelium **(Figs. 1B, 7B**, green). These cells migrate from the dorsal neural tube and populate pharyngeal arches as early as E8.5 ^47^. Then, NC-derived cells closest to the pharyngeal arch endothelium differentiate into vascular smooth muscle cells, a process which is indispensable for remodeling the PAAs into the AAAs^10,14,48^.

**Figure 7.**
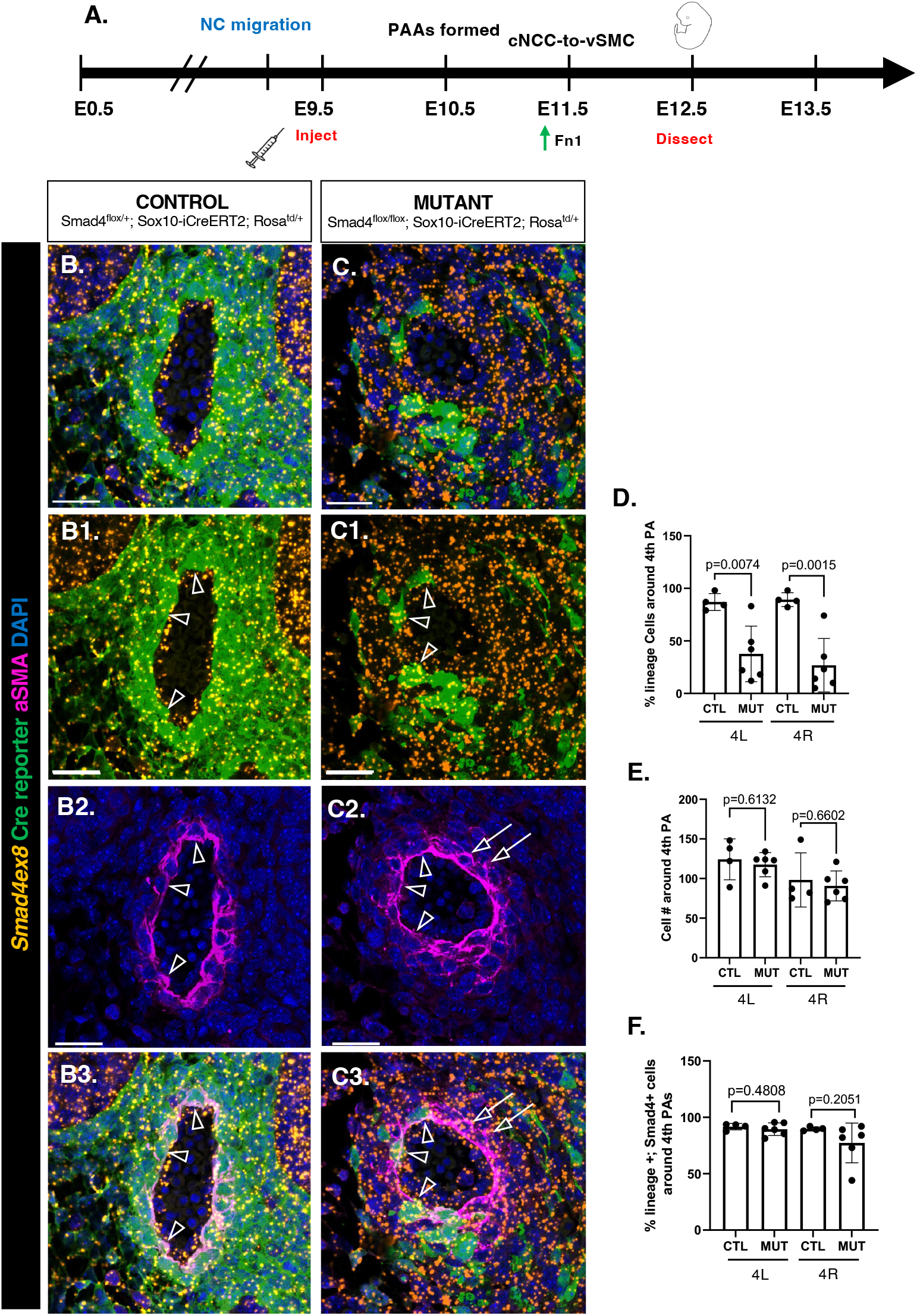
SMAD4 is required for neural crest contribution to the pharyngeal arches. Tamoxifen injection conditions (**A**). Fluorescent *in situ* hybridization for *SMAD4* exon 8 mRNA and immunofluorescence for the Cre reporter and αSMA performed on E12.5 coronal sections through the PAs of control (**B-B3**) and mutants (**C-C3**). Arrowheads point to Cre+; SMAD4+ cells in the PAs that also express αSMA. Arrows in **C2-C3** point to lineage-negative cells, which express αSMA. Percent lineage cells around the 4th PAAs within 4 cell layers closest to the endothelium is quantified in (**D**). Total cell numbers in the same region are quantified in (**E**). Percent lineage cells expressing SMAD4 is quantified in (**F**). N=4 Controls and 6 Mutants. Each dot is an average of all sections analyzed per embryo. For statistics, 2-tailed, unpaired, Student’s t-test was used; p <0.05 is considered statistically significant. Scale bars are 50 µm.

To investigate a cell-autonomous role for SMAD4 in the induction of Fn1 and αSMA expression in NCs, we used the Sox10-iCreER^T2^ strain ^49^ to conditionally ablate SMAD4 in NCs at E9.5 **(Fig. 7A)**. Sox10 is expressed in post-migratory NCs ^50,51^. Thus, by injecting tamoxifen at E9.5, we hoped to delete SMAD4 in the pharyngeal NC without causing lethality due to early deletion of SMAD4, as seen in studies using the Wnt1-Cre1 transgenic strain ^28,42,52^. Upon evaluating SMAD4 knockdown efficiency at E12.5, we made two observations. *First*, when comparing controls **(Fig. 7B-B1)** to mutants **(Fig. 7C-C1)**, we saw fewer Sox10-lineage cells in the 4^th^ pharyngeal arches **(Fig. 7D)**. However, despite the reduced number of Sox10-lineage cells around the 4^th^ PAAs in Smad4^flox/flox^;Sox10-iCreER^T2^mutants, the total number of cells surrounding the arch artery endothelium was not affected **(Fig. 7E)**. *Second*, the few Sox10-lineage cells remaining in the Smad4^flox/flox^;Sox10-iCreER^T2^ mutants still expressed *SMAD4* mRNA **(Fig. 7C-C1** and **7F)**. Moreover, while controls exhibited a fully Sox10-lineage-derived αSMA layer **(Fig. 7B2-B3)**, in mutants, primarily non-Sox10-lineage compensatory cells contributed to αSMA coverage around the endothelium **(Fig. 7C2-C3)**.

Since most of the remaining NCs in the pharyngeal arches of mutants expressed *SMAD4* mRNA, we sought to determine whether SMAD4 was ablated in other NC-derived lineages in the Smad4^flox/flox^; Sox10-iCreER^T2^ mutants. Thus, we evaluated *SMAD4* deletion in the dorsal root ganglia (DRGs), which are neural-crest-derived ^52^. Our analyses demonstrated that, on average, 97% of NC-derived cells in control DRGs expressed *SMAD4* **(Fig. 8A-A’’, C)**, while the majority of NC-derived cells in the mutant DRGs lacked *SMAD4* **(Fig. 8B-B’’, C)**. We also observed that NC-derived neurons in mutants also lacked *SMAD4* **(Fig. 8 D-E’’)**, showing that Sox10-iCreER^T2^ efficiently mediates recombination at the Rosa reporter and *SMAD4* loci. Together, these data indicate successful attenuation of SMAD4 from the Sox10 lineage, and that the expression of *SMAD4* is required for NC-derived cells to contribute to the mesenchyme of the 4^th^ pharyngeal arches, in particular.

**Figure 8.**
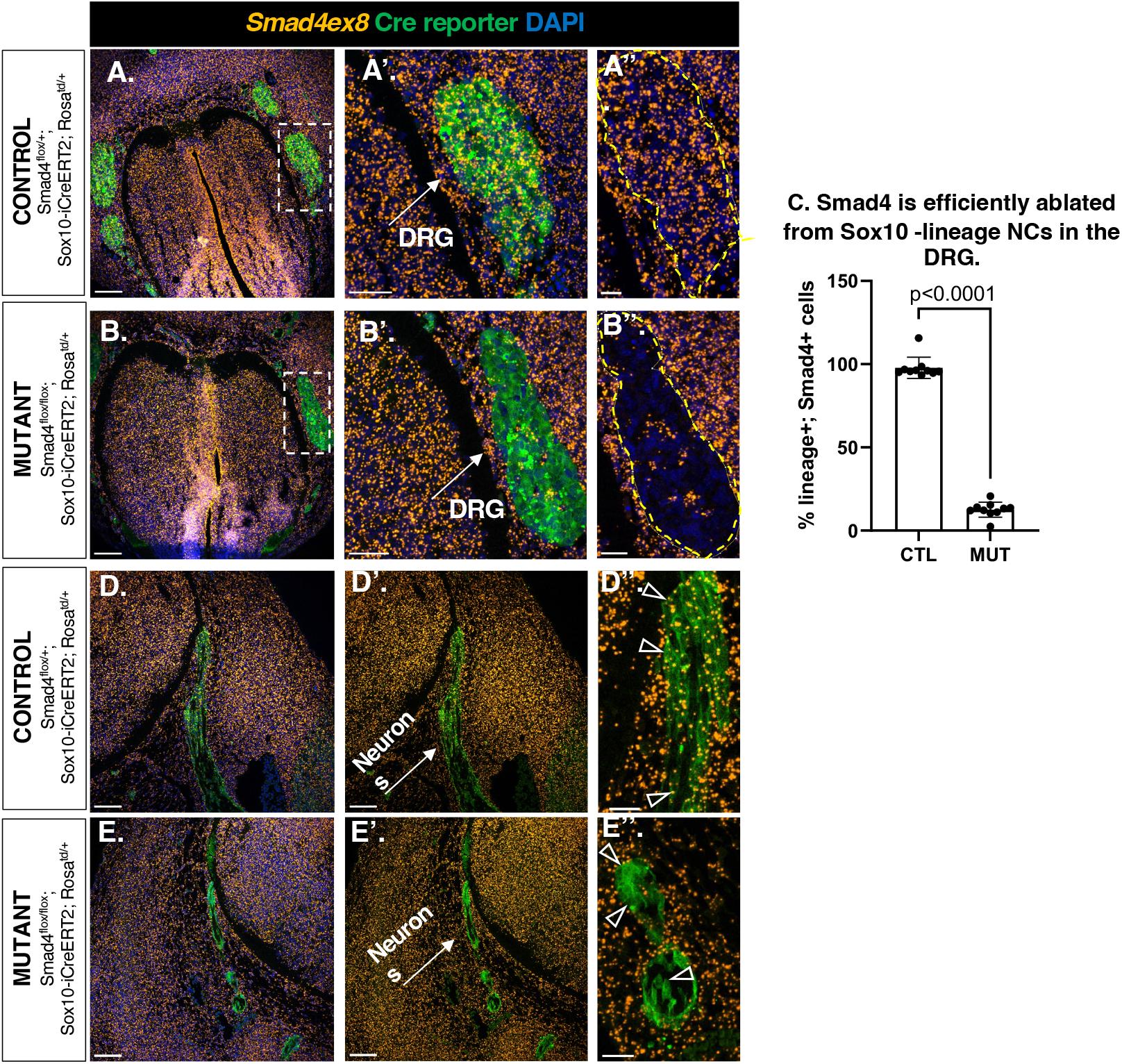
Sox10-iCreER^T2^ mediates efficient SMAD4 ablation in NC-lineage cells outside of pharyngeal arches. Images of fluorescent *in situ* hybridization for SMAD4 exon 8 mRNA and immunofluorescence for the Cre reporter on E12.5 coronal sections through the dorsal root ganglia (DRG) of a control (**A-A’’**) and mutant (**B-B’’**) embryo. Quantifications of percentage of SMAD4+ lineage cells in the DRGs (**C**), N=5 controls and N=8 mutants. Each dot is an individual section analyzed. SMAD4 is efficiently ablated in Sox10-lineage neurons (**D-E”**). Controls (**D-D’’**) and mutants (**E-E’’**). For statistics, 2-tailed, unpaired, Student’s t-test was used; p <0.05 is considered statistically significant. Scale bars in **A-B, B-B’** are 100 µm and in **A’’** and **B’’** are 50 µm.

To address whether altered differentiation and/or migration at earlier stages contributed to the depletion of SMAD4-negative NCs in the 4^th^ pharyngeal arches at E12.5, we injected tamoxifen at E9.5 **(Fig. 9A)**. We then stained and imaged entire control and mutant E10.5 embryos (between 35-37 somites) to detect patterning and differentiation of NC-derived cells and neurons, as well as endothelial cells. NCs are known to contribute to cranial neurons and glia, and thus, we hypothesized that loss of SMAD4 could alter the migration or differentiation of cardiac NCs to Tuj1+ neurons ^53^. Tuj-1 staining revealed normal patterning of the NC-derived cranial neurons when comparing the Cre-negative control **(Fig. 9B-B’)**, Smad4^flox/+^; Sox10-iCreER^T2^ Cre-positive control **(Fig. 9C-C’)**, and Smad4^flox/flox^; Sox10-iCreER^T2^ mutant embryos (**Fig. 9D-D’)**. We next examined NC trans-differentiation to endothelial cells (ECs) (**Fig. 10**). NC-to-EC differentiation has previously been shown *in vitro*, in carotid-body-derived NCs, which express endothelial markers when cultured in defined medium ^54^. However, we did not observe any Sox10-lineage cells co-expressing VEGFR2 in the craniofacial region **(Fig. 10A1, B1)**, pharyngeal region **(Fig. 10A2, B2)** or the trunk region **(Fig. 10A3, B3)**. Finally, since NC-derived cells also give rise to the enteric nervous system ^52^, we examined NC migration toward the gut endoderm by staining for the Cre-reporter. Quantification of the percentage of Sox10-lineage cells surrounding the endoderm revealed no differences in NC accumulation between Smad4^flox/+^; Sox10-iCreER^T2^ Cre-positive controls and Smad4^flox/flox^; Sox10-iCreER^T2^ mutant embryos **(Fig. 11)**. Together, these data argue against the hypothesis that SMAD4-null NC cells are reduced in E12.5 pharyngeal arches due to their aberrant migration or differentiation into other lineages.

**Figure 9.**
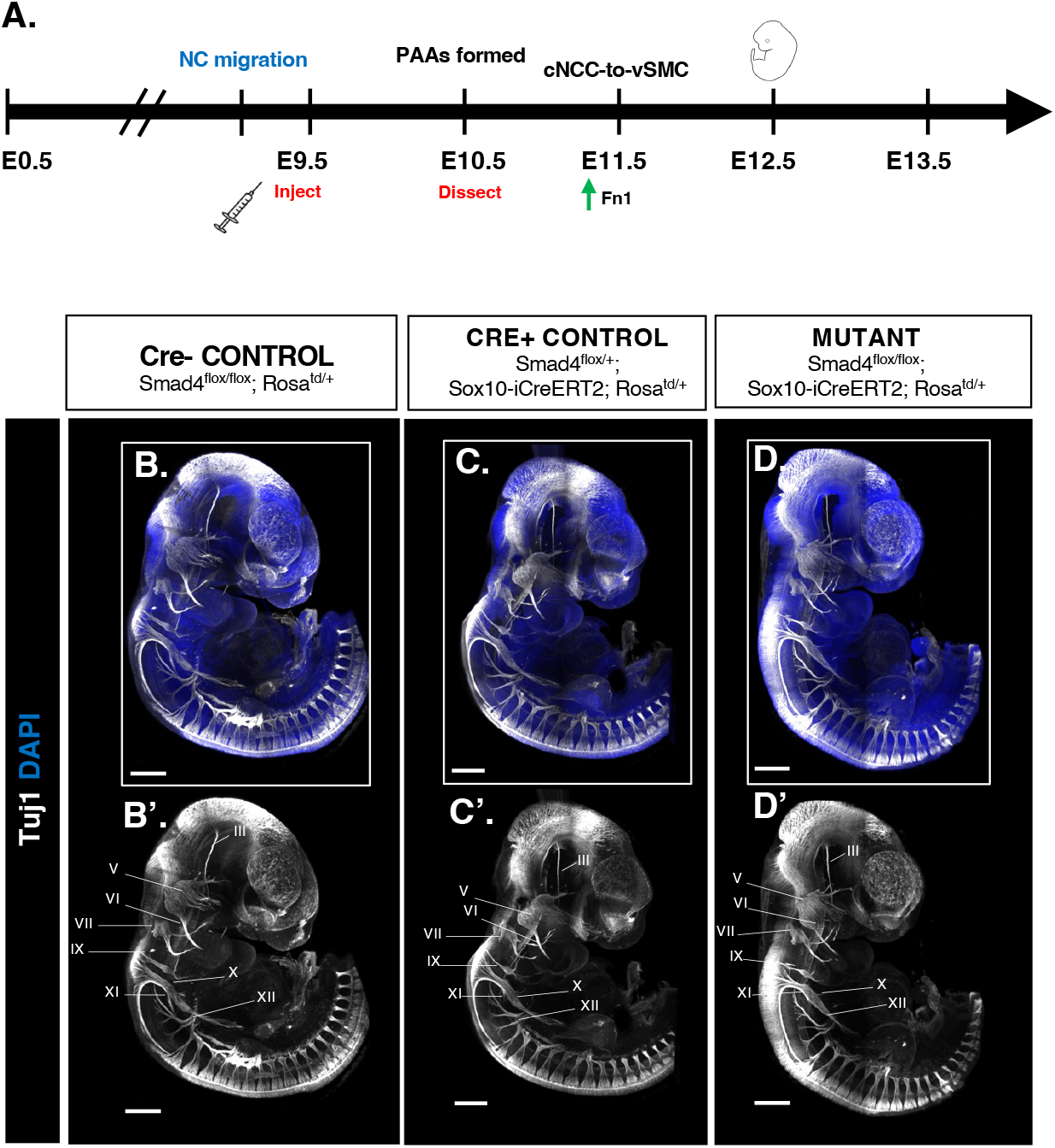
Loss of SMAD4 in the Sox10-lineage does not cause aberrant nerve patterning in E10.5 embryos. Images of whole E10.5 embryos stained for neuronal marker, Tuj1, in a Cre-negative control (**B-B’**), Cre-positive control (**C-C’**) and mutant embryo (**D-D’**). Cranial nerves are numbered with Roman numerals. Scale bars are 500 µm. Cre-negative control embryo 37 somites, Cre-positive Control embryo 37 somites, Mutant embryo 35 somites.

**Figure 10.**
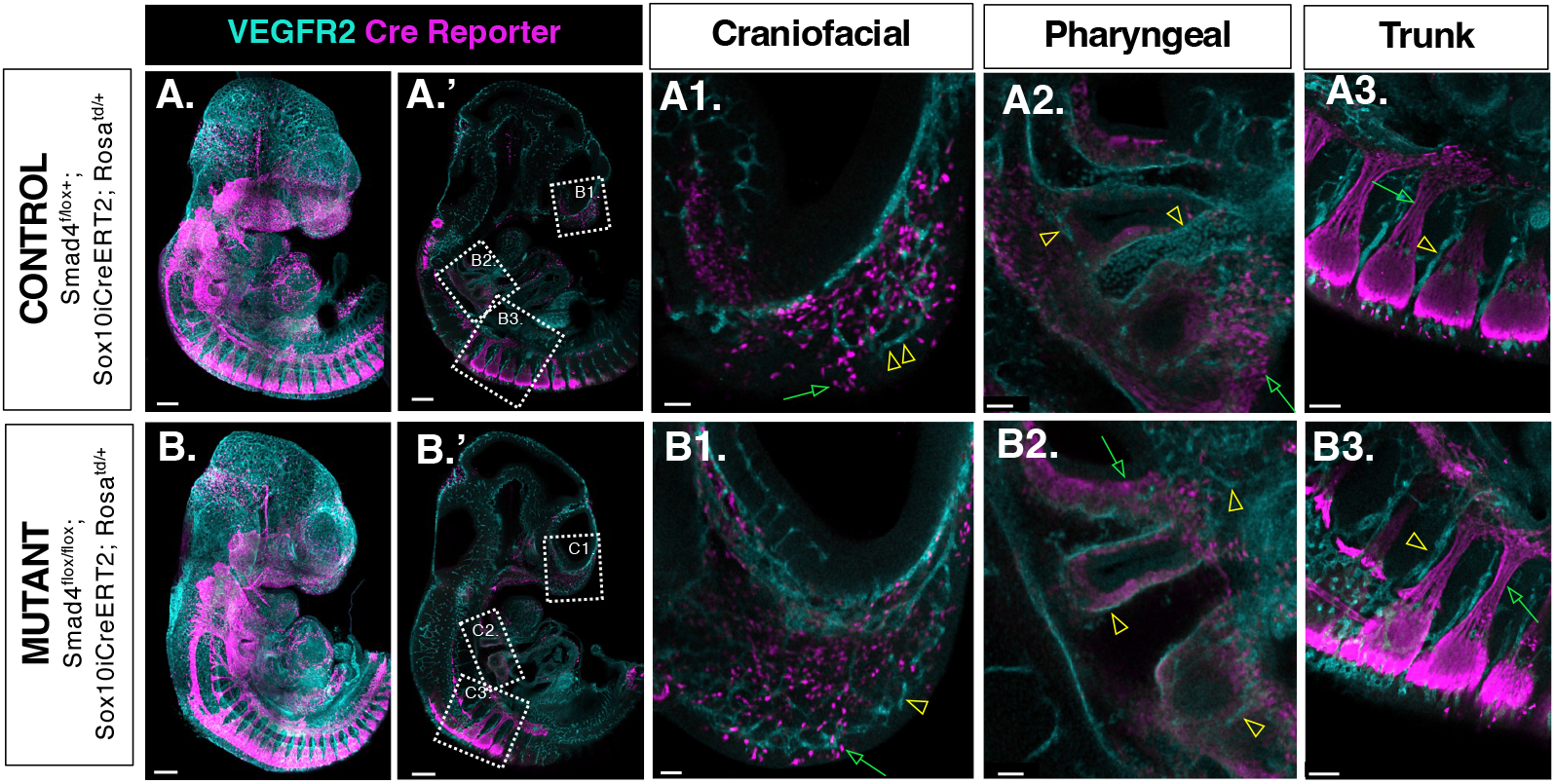
NCs do not trans-differentiate to endothelial cells in Smad4^flox/flox^;Sox10-iCreER^T2^ mutants. Images of whole E10.5 embryos stained for endothelial marker, VEGFR2, and the Cre reporter in a Cre+ control **(A-A3)** and mutant **(B-B3)** embryo. Boxed regions in **A’** and **B’** magnify the craniofacial **(A1, B1)**, pharyngeal **(A2, B2)** and trunk **(A3, B3)** regions in the control and mutant embryo, respectively. Green arrows point to VEGFR2-negative, Sox10-lineage cells and yellow arrowheads point to VEGFR2-positive, Sox10-lineage-negative endothelial cells. Scale bars in **A-A’** and **B-B’** are 300 µm and in **A1-A3, B1-B3** are 50 µm. Cre+ Control embryo 37 somites, Mutant embryo 35 somites.

### 2.4 SMAD4-null neural crest cells do not persist in the pharyngeal arches after E10.5

Previous studies using Wnt1-Cre1 transgenic mice to ablate SMAD4 have indicated that SMAD4 is important for the survival of NCs contributing to the cardiac outflow tract and pharyngeal region ^42,55^. Because we saw so few SMAD4-null NCs cells in the pharyngeal arches of E12.5 Smad4^flox/flox^; Sox10-iCreER^T2^ mutant embryos (**Fig. 7**), we hypothesized that NC survival in the pharnygeal arches was attenuated with Cre-mediated SMAD4 loss. To test this, we analyzed the presence of Sox10-lineage cells in the pharyngeal arches 30 hours after tamoxifen administration in Smad4^flox/+^;Sox10-iCreER^T2^ control and Smad4^flox/flox^;Sox10-iCreER^T2^ mutant embryos ranging from 35-37 somites (E10.5). Already at this early time after the injection of tamoxifen, we found fewer Sox10-lineage cells in the 4^th^ pharyngeal arches of Smad4^flox/flox^; Sox10-iCreER^T2^ mutants compared to Smad4^flox/+^; Sox10-iCreER^T2^ controls (**Fig. 12**). This suggests that Sox10-lineage cells in the pharyngeal arches are highly sensitive to the loss of SMAD4. Since NC-derived cells already populate pharyngeal arches at the time of injection ^37^, these studies indicate that the deletion of SMAD4 causes the loss of pharyngeal NCs, likely due to their apoptosis ^28,42,55^. Taken together, these data demonstrate the crucial role of SMAD4 in the contribution of NC-derived cells to the pharyngeal arch mesenchyme and show that relative to other NC-derived cell types, the cardiac NC is especially sensitive to SMAD4 dosage.

## 3. Discussion

SMAD4 is a common signaling mediator in the canonical TGFβ and BMP pathways. While SMAD4 has well-documented tumor suppressor activity ^24,56^, recently, missense mutations in SMAD4 have been linked to a genetic predisposition for congenital heart disease (CHD) ^22,25,26^. Because of the evolving role of SMAD4 in CHD phenotypes, we sought to further elucidate its contribution to the morphogenesis of the 4^th^ pharyngeal arch arteries, whose remodeling defects, including interrupted aortic arch, can be life-threatening if left untreated ^8,57^. To do this, we probed the role of SMAD4 in vascular smooth muscle differentiation of the cardiac neural crest, a process which is indispensable for arch artery morphogenesis.

### SMAD4 regulates αSMA expression in the 4^th^ PAs

NC differentiation to vSMCs is critical in proper arch artery morphogenesis ^10,19,58–60^. Failure of NCs to differentiate into vSMCs can result in premature PAA regression. Various factors have been implicated in vSMC differentiation of the neural crest *in vivo*, including ALK2, NOTCH signaling, Fn1, and the SMAD2/MRTFβ signaling axis ^10,31,37,61,62^. When using the R26R^CreERT2^ strain ^43^ to achieve global loss of SMAD4, we observed small (1.5-fold) and graded levels of SMAD4 deletion **(Fig. 2)**. Despite the small changes in SMAD4 expression, αSMA expression was attenuated in 60% of mutants **(Fig. 3)**. Defects in vSMC differentiation were unlikely caused by apoptosis, since the percentage of apoptotic cells, although increased in the mutants, were overall scarce (<3%) **(Fig. 3)**. These studies suggest that the role of SMAD4 in the differentiation of NC cells to smooth muscle cells is uncoupled from its role in cell survival, consistent with earlier *in-vitro* reports ^28^.

### Potential role of SMAD4 in the regulation of Fn1 expression in the 4^th^ PAs

To explore potential mechanisms underlying how SMAD4 regulates NC-to-vSMC differentiation, we investigated SMAD4 regulation of the extracellular matrix protein, Fn1. Our lab previously found that Fn1 has a dynamic expression pattern in NCs and that it is critical for NC-to-vSMC differentiation and the remodeling of the PAAs into the AAAs ^37^. There are a number of studies which indicate that TGFβ stimulation induces Fn1 synthesis both *in vitro* and *in vivo* in fibrotic tissue, where TGFβ ligands are often seen upregulated as part of the inflammatory response (Hocevar et al., 1999; Ignotzs & Massague§, 1986; Walton et al., 2017).

We examined the expression of *Fn1* mRNA and Fn1 protein in global Smad4^flox/flox^; R26R^CreERT2^ mutants, and we found that the expression of *Fn1* mRNA and protein were attenuated only in the severe mutant, while none of the mild mutants were affected **(Fig. 4)**. Thus, in contrast to αSMA, Fn1 expression was less sensitive to changes in SMAD4 levels. Additionally, these data suggested that a minimum threshold level of SMAD4 expression was sufficient for Fn1 induction around the 4th PAAs. When that threshold was met (as in mild mutant embryos), Fn1 levels were maintained. These data indicate that SMAD4 may have a role in Fn1 regulation *in vivo*, but that the regulation of αSMA expression by SMAD4 is independent of Fn1.

### Endothelial SMAD4 is not required for αSMA or Fn1 expression

It has been previously shown that Tie2-Cre mediated loss of SMAD4 from the endothelium results in embryonic lethality by E10.5, with embryos exhibiting defects in heart and vessel formation. To ablate SMAD4 in a narrow window of time and avoid the lethality associated with the early ablation of SMAD4, we utilized the Cdh5-iCreERT2 strain to delete SMAD4 from the endothelium at E10.5. The Cdh5-iCreERT2 strain allowed for efficient downregulation of SMAD4 expression in the endothelium **(Fig. 5)**. Despite the near-complete loss of SMAD4 in the PAA precursors, we did not observe anomalies in pharyngeal arch formation and growth. *Fn1* mRNA levels produced by endothelial cells were unchanged, and *Fn1* mRNA and protein expression in the surrounding NC cells were also unaffected. Similarly, there were no changes in αSMA expression in the surrounding NC-derived cells **(Fig. 6)**. Thus, our data indicate that fluctuations in αSMA and Fn1 observed in Smad4^flox/flox^; R26R^CreERT2^ mutants are attributable to SMAD4 expression changes in another cell type besides endothelial cells.

### The expression of SMAD4 in the NC is required for the contribution of NC-derived cells to the pharyngeal arch mesenchyme

To determine if SMAD4 plays a cell-autonomous role in NC-to-vSMC differentiation and to prevent NC death, we utilized the Sox10-iCreER^T2^ inducible strain to conditionally delete SMAD4 from the NC ^49^. We found a reduced number of Sox10-lineage cells surrounding the 4^th^ pharyngeal arches in mutants compared to controls **(Fig. 7)**. Additionally, despite the successful ablation of *SMAD4* in NC-derived tissues such as the dorsal root ganglia, the few Sox10-lineage (Cre+) cells remaining at E12.5 in the 4^th^ PAs expressed *SMAD4 mRNA* **(Figs. 7-8)**. This finding indicates the indispensable role of SMAD4 in the contribution of NC-derived cells to the pharyngeal mesenchyme.

Since SMAD4-null NCs were not observed in the PAs in E12.5 embryos, we sought to identify their fate. Cardiac NCs, which contribute to the developing pharyngeal arches and the cardiac outflow tract, are part of a larger population of cranial NCs ^17^. In addition to cranial placodes, cranial NCs contribute to sensory neurons as well as the sensory ganglia in cranial nerves I, II, V, VII, VIII, IX, X and XI of the peripheral nervous system ^66,67^. Thus, to determine if SMAD4-null cells take on an alternative neuronal fate, we examined the patterning of Tuj-1 expressing neurons in Smad4^flox/flox^; Sox10-iCreER^T2^ mutants compared to Smad4^flox/+^; Sox10-iCreER^T2^ Cre positive controls and Cre negative controls. We did not observe differences in the patterning of NC-derived neurons and glia in mutants compared to Cre-positive and Cre-negative controls **(Fig. 9)**. This is consistent with previous studies where Wnt1-Cre1 strain was used to ablate SMAD4, except that Ko et al., and Nie et al., observed hypoplasia of trigeminal nerve (V), which we did not detect ^41,42^.

We also examined the expression of the endothelial marker, VEGFR2, in various NC-populated regions *in vivo*, as it has been shown that NC-derived cells can differentiate into endothelial cells when cultured in defined media ^54^. When examining the craniofacial, pharnygeal and trunk regions, we did not find any Cre-reporter + Sox10-lineage cells which also expressed VEGFR2 in Smad4^flox/flox^; Sox10-iCreER^T2^ mutants, indicating that SMAD4-null NCs did not acquire an endothelial cell fate **(Fig. 10)**. Finally, we analyzed the migration of NC-derived cells to the gut, which is essential for the development of the enteric nervous system but found no difference in the percentage of NC-lineage cells surrounding the endoderm in Smad4^flox/flox^; Sox10-iCreER^T2^ mutants compared to Smad4^flox/+^; Sox10-iCreER^T2^ controls at E10.5 **(Fig. 11)**. Overall, our experiments analyzing the expression of Tuj-1, VEGFR2, and the Sox10-lineage reporter indicated that loss of SMAD4 in the NC lineage did not expand the fate potential of NCs or cause aberrant migration of these cells.

**Figure 11.**
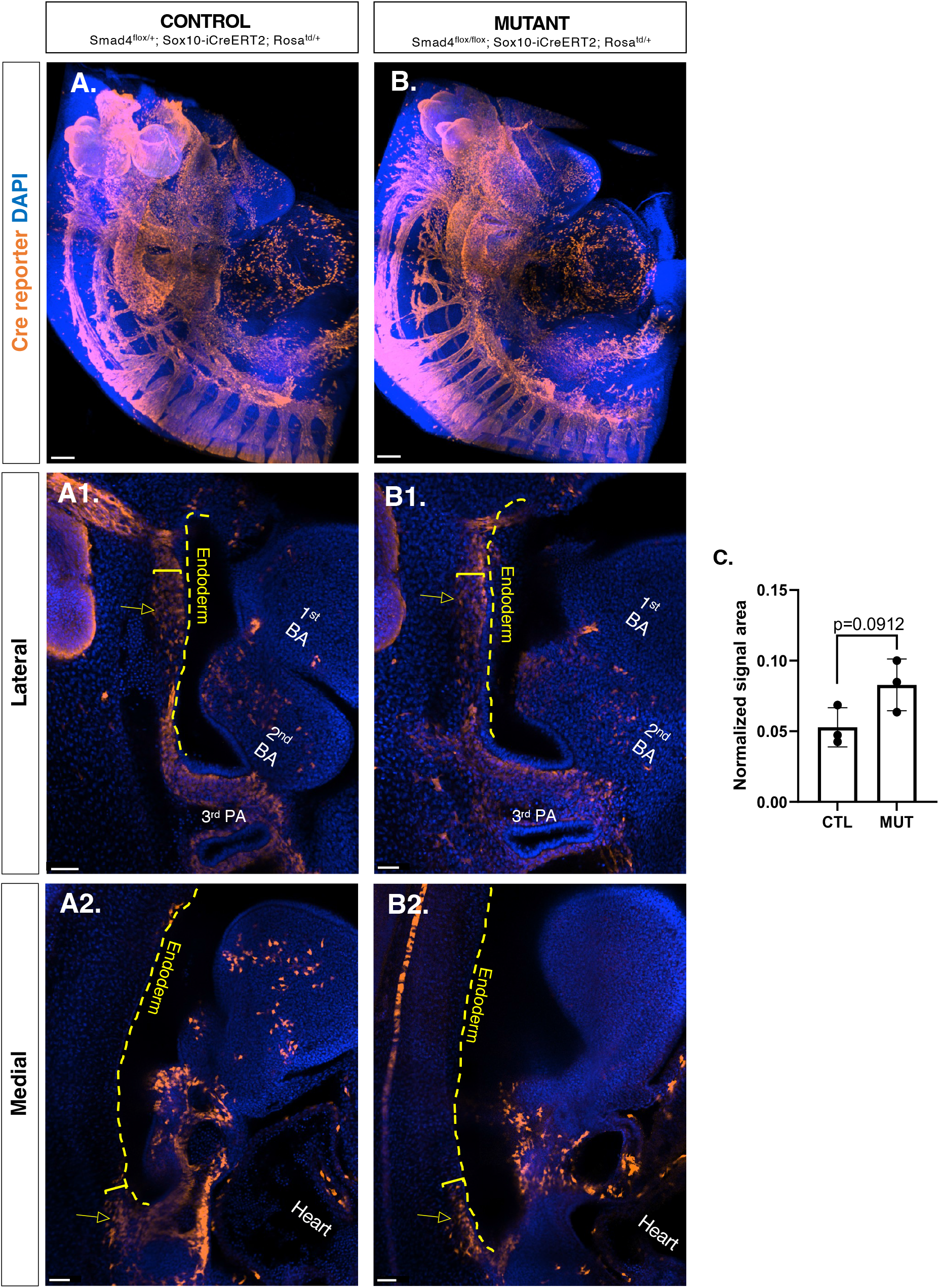
NC contribution to the enteric nervous system in Smad4^flox/flox^;Sox10-iCreER^T2^ embryos. Images of whole E10.5 embryos stained for the Cre reporter in a control **(A-A2)** and mutant **(B-B2)** embryo. Lateral **(A1, B1)** and medial **(A2, B2)** slices through the endoderm are shown. The percent of lineage cells surrounding the endoderm is quantified in **(C)**. Each dot is an average of sections analyzed for each embryo. For statistics, 2-tailed, unpaired, Student’s t-test was used; p <0.05 is considered statistically significant. N=3 control and N=3 mutants. Scale bars in **A-B** are 150 µm and in **A1-B2** are 50 µm. The embryos shown are 35 somites.

NC-derived cells are observed in the pharyngeal arches at E9.5 ^47^, the time of tamoxifen injection. Quantification of NC-derived cells at E10.5, 30 hours after the injection of tamoxifen, showed a statistically significant decrease in the number of Sox10-lineage cells in the PAs of mutants compared to controls, suggesting that the deletion of SMAD4 in these cells resulted in their death **(Fig. 12)**. This finding demonstrates a stringent requirement for SMAD4 for the contribution of NCs to the pharyngeal arch mesenchyme.

**Figure 12.**
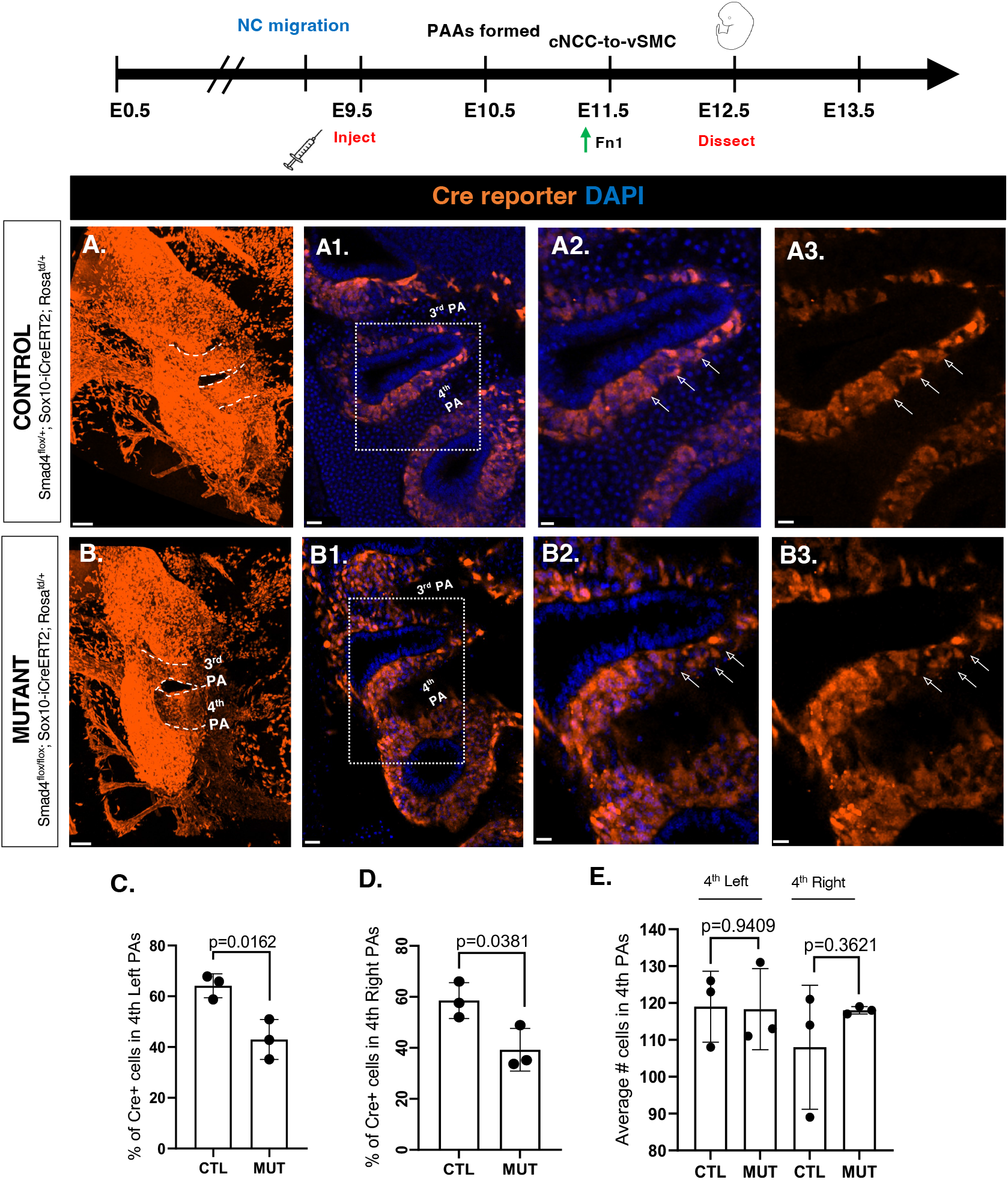
Smad4^flox/flox^; Sox10-iCreER^T2^ mutants have reduced numbers of lineage cells in the PAs at E10.5. Images of whole E10.5 embryos stained for the Cre reporter in a Cre+ control **(A-A3)** and mutant **(B-B3)** embryo. Dashed white lines in (**A, B**) outline PAs. Boxed regions in **A1** and **B1** are magnified in **A2-A3** and **B2-B3**, respectively. White arrows indicate Sox10-lineage cells in the control (**A2-A3**) and non-Sox10-lineage cells in the mutant (**B2-B3**). Percent Cre+ lineage cells in 4^th^ left and 4^th^ right PAs are quantified in **(C)** and (**D)**. Total number of cells in 4^th^ left and right PAs is quantified in **(E)**. For statistics, 2-tailed, unpaired, Student’s t-test was used; p <0.05 is considered statistically significant. N=3 control and N=3 mutants. Each dot is an average of sections analyzed for each embryo. Scale bars in **A-B** are 100 µm; scale bars in **A1-B1** are 50 µm and in **A2-B3** are 20 µm. Embryos shown are 35 somites.

Despite the reduced number of Sox10 lineage cells, the total number of cells in the 4^th^ pharyngeal arches were equivalent between controls and mutants. Moreover, our experiments indicated that the αSMA+ cell layer surrounding the PAA endothelium was mainly formed by SMAD4+ cells of non-NC origin, suggesting that cells of non-NC origin compensate for the reduced numbers of SMAD4+ NC-derived cells.

In addition to NCs, embryonic pharnygeal arches contain lateral plate mesodermal cells and thus, we hypothesize that the non-Sox10-lineage cells which contribute to the αSMA layer may be of mesodermal origin; additional studies are needed to investigate this hypothesis. It is known that mesodermal cells can give rise to the majority of vSMCs throughout the body ^68–70^. One future direction for this work would be to examine the phenotypic consequences of vSMCs formed from non-NC-derived cells compared to those that are NC-derived. While the smooth muscle layer is still formed, the function of mesodermal-derived vSMCs may differ from NC-derived vSMCs. It has been gleaned from in-vitro studies that SMCs of different origins have different phenotypic and genetic features, including cell shape, regulatory genetic network, as well as response to growth factors and inhibitors^68,69,71,72^. Furthermore, these differences may underlie vascular disease states, including aortic aneurysms and vascular calcification ^73–75^.

Overall, our studies shed light on the role of SMAD4 in NC-to-vSMC differentiation. Our data suggest that the requirement for SMAD4 in NC survival and vSMC differentiation can be uncoupled and that SMAD4 may also be a regulator of Fn1 induction in the NC. We also identified the exquisite sensitivity of cardiac NCs to SMAD4 loss, and the requirement of SMAD4 for NC contribution to the pharyngeal arch mesenchyme. Given that SMAD4 is becoming an increasingly important target gene in CHD pathology, these studies offer additional insight into its functional role during arch artery morphogenesis.

## 4. Experimental Procedures

### 4.1 Mouse strains and Generation of Mutants

All animals were maintained in accordance with the regulations of Rutgers Animal Care and the Rutgers International Animal Care and Use Committee (IACUC). The following mouse strains were used in this study (see **Table 1**): Smad4^tm2.1cxd^/J ^45^; JAX Stock no. 017462, here referred to as **Smad4^flox/flox^**, B6.129-Gt(ROSA)26Sortm1(cre/ERT2)Tyj/J ^43^; JAX Stock no. 008463, here referred to as **R26R^CreERT2/CreERT2^**, CBA;B6-Tg(Sox10-icre/ERT2)388Wdr/J ^49^; JAX Stock no. 027651, here referred to as **Sox10-iCreER^T2^**, and B6.Cg-Gt(ROSA)26Sortm9(CAG-tdTomato)Hze/J (^76^); JAX Stock no. 007909, here referred to as **R26R^Td/Td^** mice.

**Table 1:** Detailed list of commercially available strains used in this study.

| Strain Name | Stock No. | Reference | Manuscript shorthand |
| --- | --- | --- | --- |
| Smad4<sup>tm2.1cxd/J</sup> | **017462** | Yang et al., 2002 | Smad4<sup>flox/flox</sup> |
| B6.129-Gt(ROSA)26Sortm1(cre/ERT2)Tyj/J | **008463** | Ventura et al., 2007 | R26R<sup>CreERT2</sup> |
| CBA;B6-Tg(Sox10-icre/ERT2)388Wdr/J | **027651** | McKenzie et al., 2014 | Sox10-iCreER<sup>T2</sup> |
| B6.Cg-Gt(ROSA)26Sortm9(CAG-tdTomato)Hze/J | **007909** | Madisen et al., 2010 | R26R<sup>Td/Td</sup> |

C57BL/6-Tg(Cdh5-Cre/ERT2)1Rha mice ^77^ were a gift from Dr. Ralf Adams, Max Planck Institute for Molecular Biomedicine, MÜnster, Germany.

Smad4^flox/flox^;R26R^Td/Td^ females were generated by crossing Smad4^flox/flox^ and R26R^Td/Td^ mice. Smad4^flox/+^;R26R^CreERT2/TdTom^ males were generated by crossing Smad4^flox/flox^;R26R^Td/Td^ females to R26R^CreERT2/CreERT2^ males. Smad4^flox/+^;Sox10-iCreER^T2^ males were generated by crossing Smad4^flox/flox^;R26R^Td/Td^ females to Sox10**-**iCreER^T2^ males. To generate Smad4^flox/+^;Cdh5-CreERT2 males, Smad4^flox/flox^;R26R^Td/Td^ females were crossed with Cdh5-CreERT2 males.

To ablate SMAD4 globally, Smad4^flox/flox^; R26R^Td/Td^ females were crossed with Smad4^flox/+^;R26R^CreERT2/Td^ males. To ablate SMAD4 in the endothelium, Smad4^flox/flox^; R26R^Td/Td^ females were crossed with Smad4^flox/+^;Cdh5-CreERT2 males, and to ablate SMAD4 in the neural crest, Smad4^flox/flox^; R26R^Td/Td^ females were crossed with Smad4^flox/+^;Sox10-iCreERT2 males. Pregnant mothers were injected intraperitoneally with Tamoxifen to induce Cre-Lox recombination, as described below.

### 4.2 Tamoxifen Injections

Tamoxifen (Sigma Aldrich, T5648) was dissolved in sesame oil (Sigma Aldrich, S3547) at a concentration of 10 mg/ml via vortexing for 2 hours at room temperature before intraperitoneal administration into pregnant mothers. For embryos harvested 24-30 hours post-injection, 1.5 mg of tamoxifen was injected at 10 am of E9.5. For embryos harvested 56 hours post-injection, 3 mg of tamoxifen dissolved in 300 µl of corn oil (MP Biomedicals, 901414) was injected at 8 am of E10.5.

### 4.3 Genotyping

To obtain embryonic DNA, murine embryonic yolk sacs were incubated overnight in lysis buffer (1M Tris pH 8.5; 0.5M EDTA; 5M NaCl; 20% SDS) containing 0.02 mg/ml proteinase K (Thermofisher, EO0492) at 58° C in a ThermoMixer F1.5 (Eppendorf). To precipitate DNA, 100% isopropanol was added to each sample, and precipitated DNA was dissolved in dH_2_O overnight at 37° C in a ThermoMixer. Genotyping was done by PCR. For genotyping the SMAD4 floxed allele, primers listed in ^28^ were used, resulting in a 282 bp PCR product for the wild-type allele and a 330 bp PCR product for the floxed allele. For detecting the R26R^CreERT2^ allele, Cre-specific primers 5’-CTA GAG CCT GTT TTG CAC GTT C-3’ and 5’-GTT CGC AAG AAC CTG ATG GAC-3’ were used, resulting in a 320 bp PCR product. For detecting the Sox10-iCreER^T2^ allele, the following primers to the *improved* Cre (iCre) were used: 5’-CTG TGG ATG CCA CCT CTG ATG-3’ and 5’-GCC AGG TTC CTG ATG TCC TG-3’ generating a 442 bp PCR product.

### 4.4 Tissue-processing and preparation of formalin-fixed paraffin embedded (FFPE) sections

Embryos were dissected in cold 1X Phosphate-buffered saline (PBS), prepared from a 10% stock solution (Thermofisher, J75889-K2), and fixed in 10% Neutral-buffered Formalin (VWR, 10790-714) for 16 hours at room temperature. Embryos were then washed in 1X PBS and dehydrated via a series of graded ethanol (EtOH) solutions made in water. For E12.5, embryos were incubated for 1 hour at room temperature in 70%, 80%, and then 90% EtOH with agitation. Embryos were then incubated twice in 100% EtOH, for 1 hour each, at room temperature. Embryos were then incubated in Xylene twice for 30 min each, and in paraffin 70° C, twice for 30 min each. For E10.5, embryos were incubated in 70% EtOH for 3 min, and 80% EtOH for 3 min. Then, embryos were incubated in 95% EtOH twice, for 5 min each, then once for 10 min, and finally in 100% EtOH three times, for 10 min each. Embryos were then incubated in Xylene twice for 20 min each, and in paraffin at 70° C, twice for 20 min each. Embryos were then embedded into paraffin and 5 µm sections were collected using Leica Biosystems Rotary Microtome Manual HistoCore BIOCUT.

### 4.5 Multiplex Fluorescence in-situ hybridization

For detection of *Fn1* mRNA, tissue sections were first embedded into paraffin and cut into 5 µm sections, as described above. *In situ* hybridization was performed to detect *Fn1* mRNA using a probe (ACD, 408181) according to instructions in the Multiplex Fluorescent Kit v2 (323100) from ACD (**Table 2**, protocol **#1**). Bound probes were detected using TSA Plus fluorophores Cyanine 5 (Akoya Bioscience, NEL745001KT) and Cyanine 3 (Akoya Bioscience, NEL744001KT) according to manufacturer protocols; see **Table 2**, protocol **#2**.

**Table 2.** Links to external protocols referenced.

| Prot. # | Prot. name | Link |
| --- | --- | --- |
| 1 | Fluorescent Multiplex | https://acdbio.com/sites/default/files/UM%20323100%20Multiplex%20Fluorescent%20v2_RevB.pdf |
| 2 | Multiplex + IHC | https://acdbio.com/sites/default/files/323100-TN%20Multiplex%20Fluorescent%20V2%20with%20IF_incOpal780.pdf |
| 3 | Basescope RED + IHC | https://acdbio.showpad.com/share/3eLlrIP9sGPnD5pYCshq2 |

Protocols listed in **Table 2** were used without modifications.

Following transcript detection, immunofluorescence was performed according to the *ACD Inc*. protocol listed below (**Table 2**, protocol **#2**). The following antibodies were used: Rabbit anti-mCherry (1:1000, Abcam, ab167453), recombinant rabbit anti-Fibronectin (1:1000, Abcam, ab19056), and mouse anti-αSMA (1:500, Sigma Aldrich, A5228). Primary antibodies were detected with secondary antibodies from Invitrogen (1:300): donkey anti-rabbit conjugated to Alexa-488 (A21206) and donkey anti-mouse conjugated to Alexa-555 (A31570). Secondary antibody solutions also included DAPI (1:1000 dilution of 5mg/mL stock made in dH_2_O, Thermofisher, D3571). Following immunohistochemistry, samples were mounted with ProLong gold antifade solution (Thermofisher, P36930).

### 4.6 Base-scope in situ hybridization and Immunofluorescence

In the Smad4^flox/flox^ strain, Smad4 exon 8 is flanked by two loxP sites. Cre-mediated recombination of Smad4 exon 8 results in a null allele ^45^. Therefore, using formalin-fixed paraffin embedded sections (prepared as above) we assayed the presence of the SMAD4 exon 8 sequence (51 bps) and performed downstream immunofluorescence for the Cre reporter according to the *Basescope RED* + IHC protocol, see **Table 2**, protocol **#3** (*ACD Inc.,* 323900). Samples were mounted with ProLong gold antifade solution.

### 4.7 Whole-mount immunofluorescence

Whole-embryo immunofluorescence of E10.5 embryos was performed as previously described in ^78^. The primary antibodies used were: Rabbit anti-mCherry (1:1000, Abcam, ab167453), Mouse anti-Tuj1 (1:150, Covance, MMS-435P), and Goat anti-VEGFR2 (1:200, R&D Systems Inc., AF644). Primary antibodies were detected with secondary antibodies from Invitrogen (1:300): donkey anti-rabbit conjugated to Alexa 555 (A31572), donkey anti-mouse conjugated to Alexa-647 (A31571), and donkey anti-goat conjugated to Alexa-488 (A11055). Secondary antibody solutions also included DAPI, as above.

### 4.8 Image Visualization and Analysis

Confocal images were acquired with the Nikon A1 confocal microscope using a 20X CFI Apo LWD Lamda S water immersion objective (MRD77200). Images of sections and whole embryos were analyzed with ImageJ software (National Institutes of Health) and Imaris Viewer, version 9.9.1 (Oxford Instruments), respectively. To assay the expression of Smad4 in the pharyngeal region of R26R^CreERT2^ samples, a defined region of interest (ROI) was drawn within the pharyngeal region, and the number of Smad4 puncta per ROI was counted. Identically sized ROIs were used for analyses of control and mutant samples. To validate Smad4 loss in the endothelium and neural crest, the percentage of Smad4+ nuclei was quantified in Cre-reporter+ cells. To measure *Fn1 mRNA*, Fn1 protein, and αSMA expression levels around the pharyngeal arches, mean fluorescence intensity was measured as shown in the figures and the signal intensity was normalized to DAPI for the corresponding ROI.

## Figure Legends

**Supplementary Figure 1.**
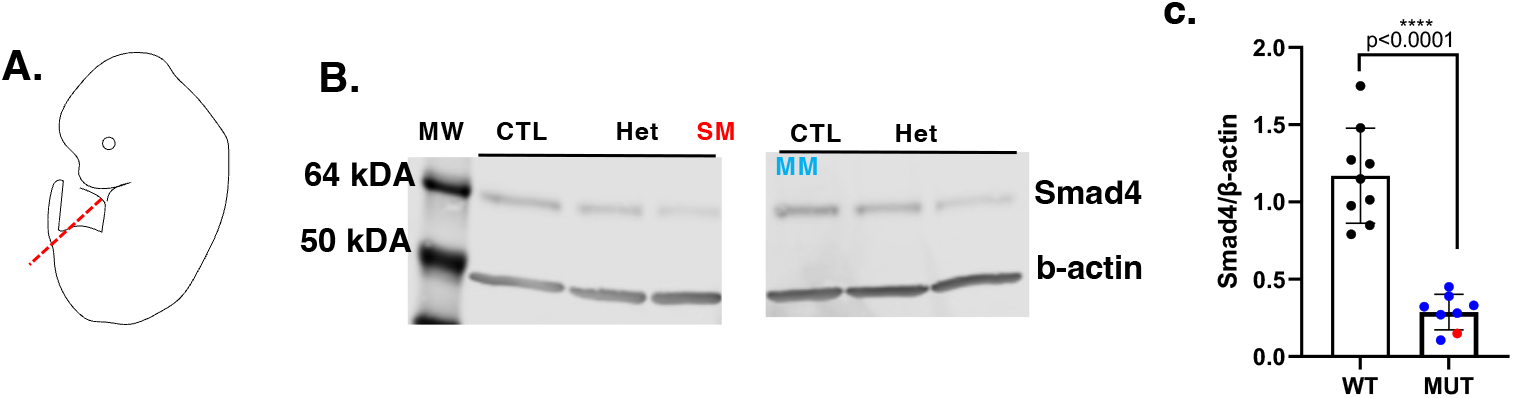
Schematic of posterior embryo region used for Western blot assays (**A**). Representative Western blot of SMAD4 proteins in the severe mutant (SM, red), mild mutant (MM, blue) Cre-positive controls (Het) and Cre-negative controls (CTL) (**B**). Quantifications of normalized SMAD4 expression levels in 9 controls and 9 mutants analyzed **(C)**.

**Supplemental Table 1.** The frequency of each genotype in harvested E12.5 embryos.

| Cross<br>Smad4<sup>flox/flox</sup>; R26R<sup>td/td</sup><br>X<br>Smad4<sup>flox/+</sup>; R26R<sup>CreERT2</sup> |  | Control<br>Smad4<sup>flox/flox</sup>; R26R<sup>td/+</sup><br>OR<br>Smad4<sup>flox/+</sup>; R26R<sup>td/+</sup> | Heterozygous<br>Smad4<sup>flox/+</sup>; R26R<sup>td/CreERT2</sup> | Mutant<br>Smad4<sup>flox/flox</sup>; R26R<sup>td/CreERT2</sup> | Total<br>N | χ<sup>2</sup> |
| --- | --- | --- | --- | --- | --- | --- |
| Observed | E12.5 | 79 (48.2%) | 38 (23.2%) | *47 (28.6%) | 164 | 0.614 |
| Expected |  | 82 (50%) | 41 (25%) | 41 (25%) |  |  |

## Acknowledgments

We would like to thank all members of the Astrof lab for their careful editing of this manuscript: Michael Warkala, Dr. Cecilia Arriagada and Dr. Gideon Obeng. We would also like to thank Drs. Melissa Rogers, Steven Levison and Theresa Wood of the Rutgers Biomedical Sciences Graduate Program for their feedback with experiments. We would also like to thank the faculty and staff of the Rutgers University Animal Care Facilities for assistance with animal husbandry. BEA would also like to thank Sam Russo for his help with tissue histology, Michael Warkala and AnnJosette Ramirez for their help with embryo handling and Shabazz Hall for her support and encouragement in the preparation of this manuscript.

